# Signal integration by a bHLH circuit enables fate choice in neural stem cells

**DOI:** 10.1101/2022.10.10.511605

**Authors:** Nagarajan Nandagopal, Alexsandra Terrio, Fernando Z. Vicente, Ashwini Jambhekar, Galit Lahav

## Abstract

Stem cells integrate information from multiple signals in their environment to make fate decisions. It is unclear how signal integration is linked to the coordinated activation of a target fate program and simultaneous inactivation of competing fates. Here, we investigated this question in mouse neural stem cells, which differentiate synergistically into astrocytes in response to combined treatment with Bone Morphogenetic Protein (BMP) and Leukemia Inhibitory Factor (LIF) at the expense of alternative neuronal or oligodendrocyte fates. Analysis of the expression dynamics of Glial Fibrillary Acidic Protein (GFAP), an early astrocyte marker, showed that its synergistic activation in BMP and LIF reflects early activation by LIF which is sustained by a delayed contribution to its expression from BMP. In parallel, multiplexed RNA-FISH analysis of 14 basic helix-loop-helix (bHLH) transcription factors, known to regulate alternative fates, showed that LIF and BMP individually control different subsets of bHLHs, but together suppress all bHLHs known to promote alternative fates. Ectopic expression experiments showed that these bHLHs also inhibit GFAP induction, suggesting that suppression of alternative fates by BMP + LIF simultaneously relieves GFAP inhibition. In particular, BMP primarily affected inhibitory bHLHs indirectly, through induction of Id factors, explaining why it has a delayed contribution to GFAP transcription compared to LIF. These results show that a circuit of bHLH factors enables both synergistic astrocytic differentiation and suppression of alternative fates in NSCs. Signal integration by bHLH circuits for fate choice could be broadly relevant, given the widespread utilization of these and other bHLH factors across diverse developmental contexts.

## Introduction

Cells can combine information from multiple signals to enact specific behaviors. This is particularly evident in the fate decisions of multipotent stem cells, which choose between different fate programs based on the specific combinations of extracellular signals they receive [1,2]. Such signal-specific fate choices involve two activities: first, integration of information from distinct signaling pathways, such that their combined effect differs from the effect of each separately, and second, coordination between fates such that genes corresponding to the target fate program are activated while alternate fate programs are suppressed. Importantly, these two activities must be coupled. How and where in the cell are signal integration and fate choice linked?

We investigated this question using neural stem cells (NSCs), which self-renew in the presence of growth factors, but can be induced to differentiate into neurons, astrocytes, or oligodendrocytes by treatment with particular combinations of chemicals or signaling ligands [3,4]. For instance, previous work has shown that treatment of NSCs with Bone Morphogenetic Protein (BMP) and Leukemia Inhibitory Factor (LIF) synergistically induces astrocytic differentiation, based on expression of an early marker of astrocyte fate, Glial Fibrillary Acidic Protein (GFAP) [5].

Integration of LIF and BMP signaling was suggested to occur through an indirect interaction between their respective downstream nuclear effectors, Stat3 and Smad1, at the GFAP promoter. However, the decision to differentiate into astrocytes involves not just activation of astrocyte-specific genes, but also suppression of neuronal and oligodendrocytic fate programs. The link between synergistic activation of the astrocyte program and suppression of alternate programs remains unclear.

Basic helix-loop-helix (bHLH) transcription factors play central roles in fate regulation in neural stem cells. bHLHs comprise a large family of transcription factors characterized by the basic helix-loop-helix domain, which mediates dimerization [6]. They are categorized into seven phylogenetic classes. Previous work has shown that the Class II bHLHs Ascl1 and Olig1/2 drive neuronal and oligodendrocytic programs, respectively [7–11]. Class II bHLHs heterodimerize with Class I bHLHs such as Tcf3, Tcf4 and Tcf12 to carry out their activities [12–15]. On the other hand, they are typically antagonized by Class V and Class VI factors - Id, Hes, and Hey - through heterodimerization or transcriptional inhibition [14–16]. Given these roles, we reasoned that the signals that induce astrocyte differentiation must also signal to bHLHs that control alternative fates.

Here, we show that LIF and BMP both synergistically activate GFAP and coordinate between fates through a circuit of bHLHs. We first analyzed the dynamics of GFAP activation to identify the timescale over which signal integration occurs, and then how LIF and BMP signaling impact bHLH expression on this timescale. We show that LIF and BMP rapidly regulate all bHLHs analyzed, but without apparent synergy. Despite lack of synergy, bHLH expression in individual cells has the power to predict GFAP status of cells. In fact, multiple bHLHs that promoted alternative fates also inhibited GFAP expression. Importantly, combined LIF and BMP treatment was required to downregulate or inhibit all these bHLHs, thus simultaneously relieving GFAP inhibition and suppressing alternate fates.

## Results

### Synergistic expression of GFAP reflects early activation by LIF that is sustained by a delayed contribution from BMP

NSCs can be induced to differentiate into astrocytes upon removal of growth factors and treatment with BMP and/or LIF (Figure 1a). Treatment with BMP or LIF induced low levels of GFAP, while treatment with both factors together led to a striking enhancement in GFAP protein levels by 24h, consistent with previous findings [5] (Figure 1b, S1a).

**Figure 1:**
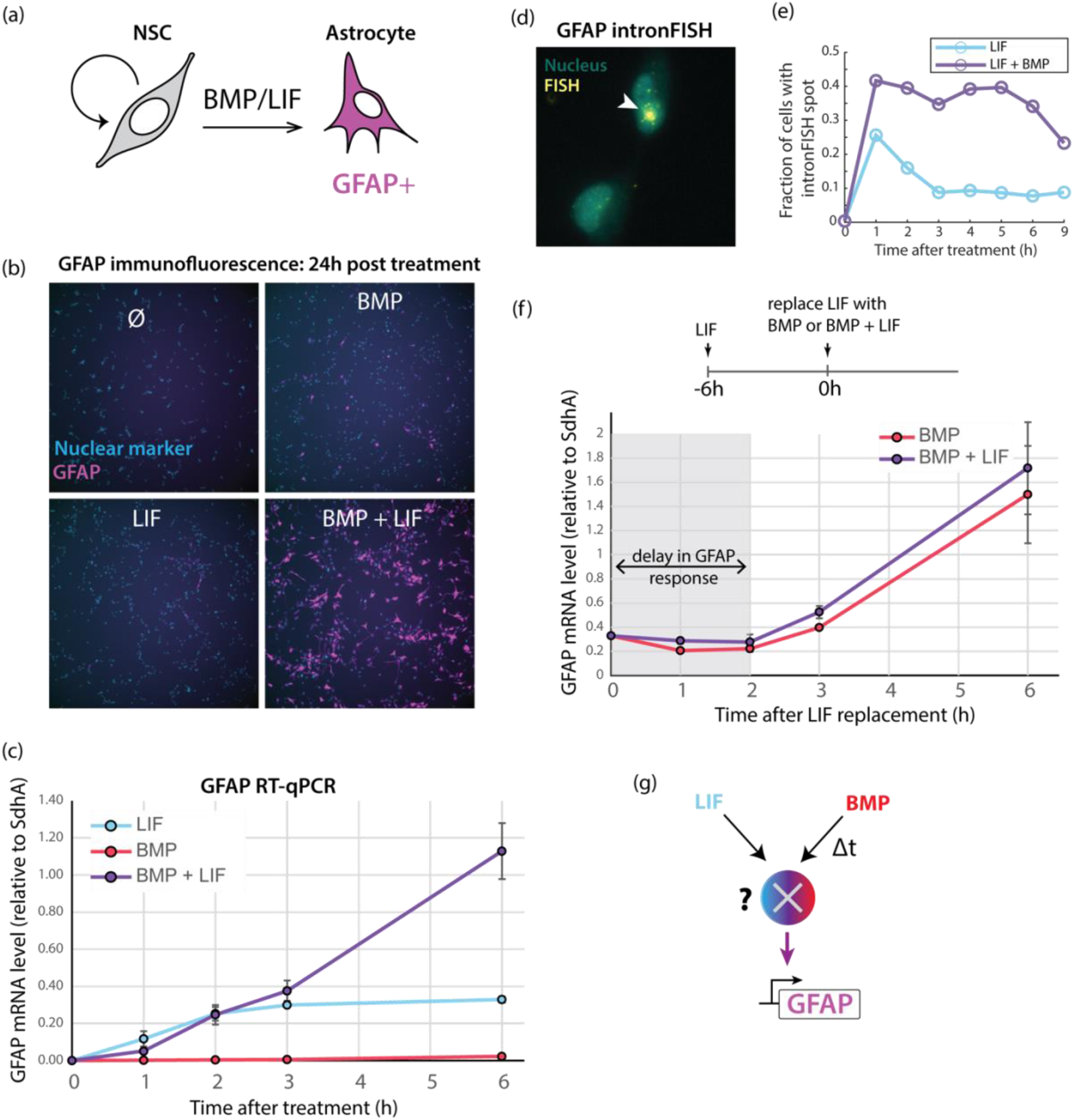
Synergy between LIF and BMP results from early activation of GFAP and its sustained transcription. (a) Mouse neural stem cells (NSCs) self-renew in the presence of EGF and FGF but differentiate into GFAP positive (‘GFAP+’) astrocytes upon treatment with BMP or LIF. (b) NSCs expressing a nuclear CFP marker (cyan) were treated with LIF, BMP or BMP + LIF for 24h or maintained in self-renewal conditions (‘Ø’) and subsequently immunostained for the differentiation marker GFAP (magenta) (c) RT-qPCR for GFAP mRNA levels over time in NSCs treated with LIF, BMP or BMP + LIF. Error bars represent S.E.M (n = 2 replicates). (d) Representative image of NSCs after HCR RNA-FISH targeting GFAP introns (‘intronFISH’) 1h hour after treatment with BMP + LIF. Active transcription of the GFAP gene results in strong localized signal (arrowhead). (e) Fraction of cells displaying GFAP intronFISH spots at different timepoints after treatment with LIF (cyan) or BMP + LIF (purple). (f) (*Top*) Experiment schematic: cells were first treated with LIF for 6h. Subsequently, LIF was replaced with BMP or BMP + LIF (t = 0). GFAP measurements were made at different timepoints after LIF replacement. (*Bottom*) RT-qPCR analysis of GFAP mRNA levels following replacement of LIF. Error bars indicate S.E.M (n = 2 replicates). (g) (*Schematic*) LIF is integrated with a delayed contribution from BMP by an unknown mechanism (‘X’) that leads to synergistic GFAP expression.

To investigate the underlying mechanisms of signal integration between LIF and BMP, we quantified GFAP expression levels during the first few hours following treatments with the two signaling ligands, either alone or in combination. Analysis of GFAP mRNA levels using RT-qPCR showed that its expression commences within 1h in BMP + LIF and that a clear difference in GFAP levels between LIF, BMP and BMP + LIF could be observed by 3 to 6h (Figure 1c). Interestingly, LIF alone could induce GFAP expression at a comparable initial rate to BMP + LIF, though GFAP levels plateaued in this condition by 2h while they continued to rise in BMP + LIF (Figure 1c). GFAP expression thus follows different dynamics in LIF vs BMP + LIF after early activation.

What underlies these differences in dynamics? To investigate whether GFAP levels plateaued under LIF treatment because of high mRNA degradation rates, we measured mRNA half-lives after inactivating transcription with actinomycin D (Figure S1b, Methods). We calculated a half-life of ∼5h for GFAP mRNA, which is not sufficiently short to account for the plateau of its mRNA levels, and instead suggests that GFAP mRNA must be produced transiently during the first 2 hours of LIF treatment.

To directly measure the dynamics of GFAP transcription we used HCR RNA-FISH [17] to target introns in GFAP, and thus detect nascent RNA (‘intronFISH’, see Methods) [18]. intronFISH signal was bimodal in the cell population. A fraction of cells displayed bright nuclear spots, which correspond to GFAP sites that are transcriptionally active [18], while others displayed little to no signal (Figure 1d). Analysis of GFAP transcription over the first 9 hours following treatment confirmed that LIF treatment leads to an immediate, but brief, period of GFAP activation. On the other hand, BMP + LIF treatment leads to sustained GFAP transcription (Figure 1e). In addition to increasing the fraction of actively transcribing (‘GFAP-active’) cells, BMP + LIF also appeared to enhance the magnitude of transcription in GFAP-active cells, based on the higher intensity of intronFISH signal in these cells compared to those in the LIF condition (Figure S1c).

Since the predominant effects of LIF and BMP on GFAP transcription appeared to occur during different time windows (LIF < 3h, BMP > 3h), we asked whether their contributions could be separated in time. We first treated cells with LIF for 6h, allowing GFAP to reach its plateau, and subsequently replaced LIF with BMP. This led to a sharp increase in GFAP expression, reaching similar levels 6h later as that achieved in cells treated simultaneously with BMP and LIF (Figure S1d). Strikingly, the increase in GFAP under sequential treatment only began after a delay of 2h following BMP addition (Figure 1f). The same effect could be observed when LIF was replaced with BMP + LIF. These data suggest that LIF and BMP contribute to different phases of GFAP expression. In particular, the effect of BMP on GFAP includes an inherent time delay. We note that phosphorylation of Smad1 upon BMP treatment is rapid (< 1h) and comparable with or without LIF pre-treatment (Figure S1e), suggesting that the delay in GFAP response following LIF pre-treatment occurs downstream of BMP pathway activation. Finally, we assessed uniformity in GFAP expression in response to BMP + LIF using HCR RNA-FISH to detect GFAP mRNA at the single-cell level (Figure S1f). This analysis showed significant intercellular variability in GFAP mRNA and protein levels by 9h.

Together, these data show that LIF and BMP have a synergistic effect on GFAP expression, with each contributing predominantly to different temporal phases of its activation and producing a heterogeneous response. Our next goal was to identify the underlying signal integration mechanism that accounts for these features of fate regulation (Figure 1g).

### bHLH transcription factors are regulated non-synergistically by BMP and LIF to suppress alternative cell fates

The decision to commence astrocytic differentiation must be accompanied by suppression of the alternative neuronal and oligodendrocytic fates. Alternative fates in NSCs are controlled by Class II and Class I bHLHs such as Ascl1, Olig1/2 and Tcf genes. These bHLHs are typically antagonized by Hes, Hey and Id genes. To analyze whether suppression of alternative fates by BMP and LIF also occurred in a synergistic manner, we monitored the expression of 14 bHLH factors in NSCs treated with BMP, LIF, or both for 6h, when synergy in GFAP expression is apparent (Figure 1c). Importantly, these bHLHs are connected through dimerization [14] or transcriptional interactions [19], such that changes in the levels of some bHLHs could affect the levels or activities of others (Figure 2a).

**Figure 2:**
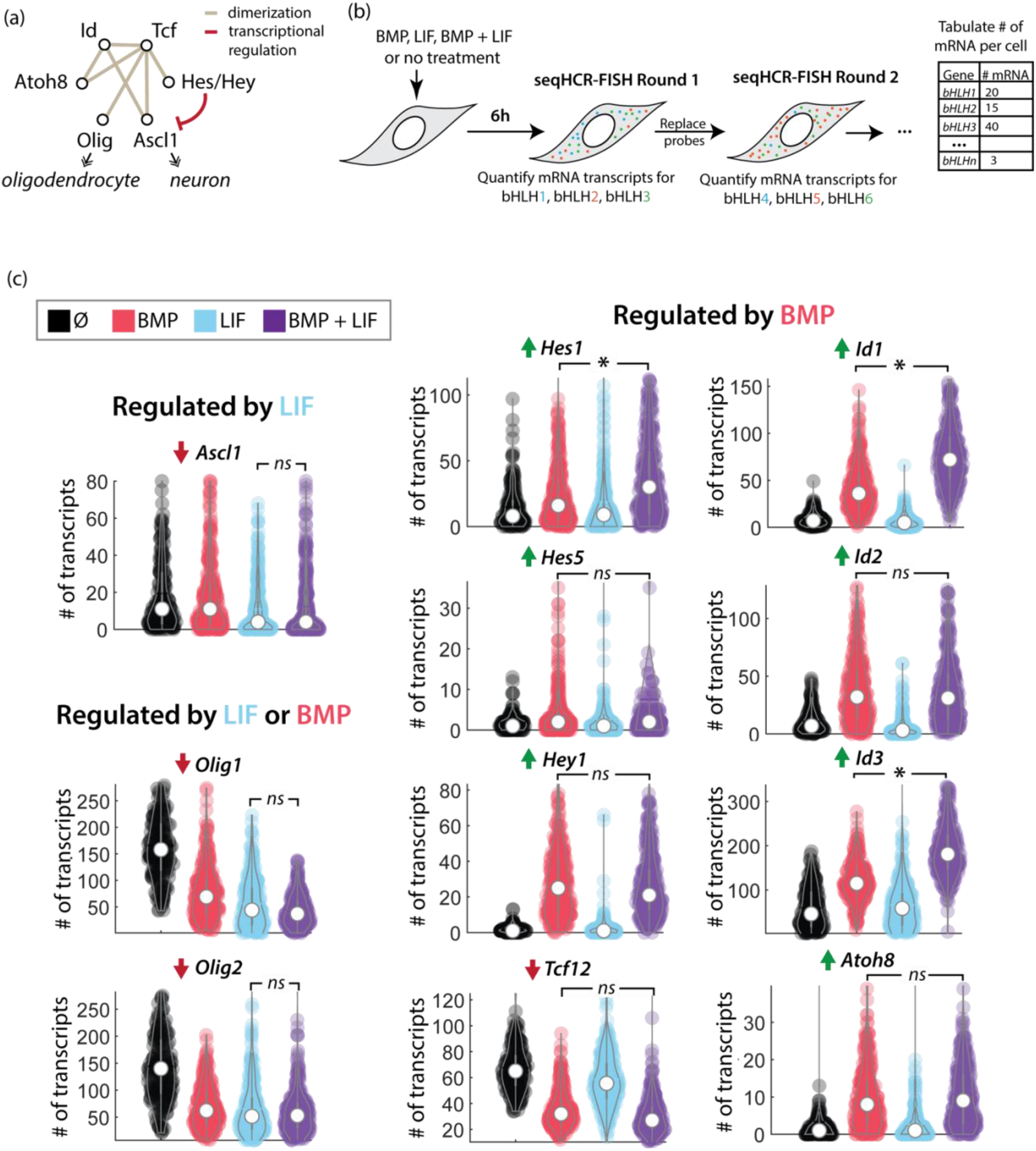
LIF and BMP regulate different subsets of bHLHs, with almost no synergy. bHLHs in neural stem cells regulate alternative neural fates and are interconnected through dimerization (brown lines) and transcriptional (red line) interactions. Note that ‘Id’ encompasses Id1, Id2, Id3, Id4, ‘Tcf’ encompasses Tcf3, Tcf4, and Tcf12, ‘Hes’ includes Hes1 and Hes5, and ‘Olig’ includes Olig1 and Olig2. (b) Schematic of experiment: NSCs were treated with different signaling ligands for 6 hours. Subsequently, sequential HCR RNA-FISH (‘seqHCR-FISH’) was used to quantify mRNA levels of multiple bHLHs in each cell over sequential rounds of mRNA detection using gene-specific probes, quantification of transcripts, and probe clearance. (d) Number of mRNA transcripts detected per cell, using sequential HCR RNA-FISH, for 11 bHLHs in self-renewal conditions (‘Ø’) or after treatment with BMP, LIF, or BMP + LIF for 6 hours. These genes display > 1.5 fold change, relative to no treatment condition, in at least one of the three signal treatments. Dark red or green arrows indicate that the gene is downregulated or upregulated, respectively, in BMP + LIF relative to no treatment condition. *‘*’* indicates a significant difference at a *P-*value of <0.01 by two-sided KS-test, while *ns* indicates no significant difference. See also Figure S2e for 3 bHLHs that show < 1.5 fold change.

We utilized sequential HCR RNA-FISH to measure all 14 bHLHs in every cell (Figure 2b, Figure S2a, Methods). This method involves sequential rounds of HCR RNA-FISH, with multiplexed detection of mRNA for up to three target genes per round. Each round is followed by DNAse treatment to eliminate FISH signal before the subsequent round. Measurements were highly reproducible between rounds and biological replicates, and FISH signal was completely cleared between rounds (Figure S2b-d). Analysis of bHLH expression 6 hours after treatment with BMP, LIF, or BMP + LIF showed that, strikingly, every bHLH responds to at least one of the two signals, with 11 out of 14 bHLHs displaying a >1.5 median fold change (Figure 2c, S2e).

Interestingly, BMP and LIF regulate different subsets of bHLHs: LIF, but not BMP, is necessary for downregulating Ascl1. On the other hand, BMP, but not LIF, regulates eight bHLHs including its canonical targets, the Id genes. Of these 8 targets, 7 are increased and one (Tcf12) is decreased. Both BMP and LIF individually downregulated Olig1 and Olig2.

Moreover, comparison of bHLH response in BMP + LIF with that in either alone showed that most (8 out of 11) responses in the combined treatment could be explained by their responses in one of the two treatments. For Hes1, Id1 and Id3, the response was moderately enhanced in BMP + LIF relative to that in BMP alone (Figure 2c).

Together, this data showed that LIF and BMP each rapidly regulate different subsets of bHLHs. In particular, both signals suppress bHLHs that drive alternative neuronal and oligodendrocytic fates. However, unlike GFAP, there is little synergy in their regulation. This implies that the suppression of alternative fates does not occur downstream of the signal integration mechanisms that lead to synergistic GFAP expression.

### bHLH expression levels in each cell has predictive power about its GFAP status

Despite displaying non-synergistic induction, we observed that bHLHs display significant cell-to-cell heterogeneity in their levels and next asked whether bHLH levels could explain the broad variability of GFAP expression observed between individual cells (Figure S1f). We investigated whether bHLH expression in individual cells could predict whether, relative to other cells, GFAP levels would be high or low. First, we measured the expression of all 14 bHLHs as well as GFAP in every cell using sequential HCR RNA-FISH every hour for 9h after treatment with BMP + LIF. We also monitored the level of LIF- and BMP-induced signaling activity in the same cells by immunostaining them for phospho-Stat3 and phospho-Smad1 following the final round of HCR RNA-FISH (Figure S3, Methods).

Next, for each analysis timepoint following BMP + LIF treatment, we trained a decision tree classifier on the expression levels of the 14 bHLHs to classify cells into GFAP-high and GFAP-low states (Figure 3a). Every trained classifier has an associated classification tree that indicates the bHLHs used as predictors in the classification (Figure 3b) and an ROC (Receiver Operating Characteristic) curve that captures the performance of the classifier across a range of decision thresholds (Figure 3c, d). Inspection of ROC curves showed that, at every analysis timepoint, bHLH levels in the cell contain information about its GFAP status. Specifically, the Area Under the Curve (AUC) for classifiers trained on bHLH levels was consistently ∼0.65, significantly greater than 0.5 expected for a random classifier and observed for classifiers trained on scrambled data (Figure 3e, Methods). Interestingly, including the levels of phospho-Stat3 and phospho-Smad1 in the classification did not significantly improve the AUC (Figure 3f), suggesting that they do not contribute additional information relevant to the GFAP status of the cell.

**Figure 3:**
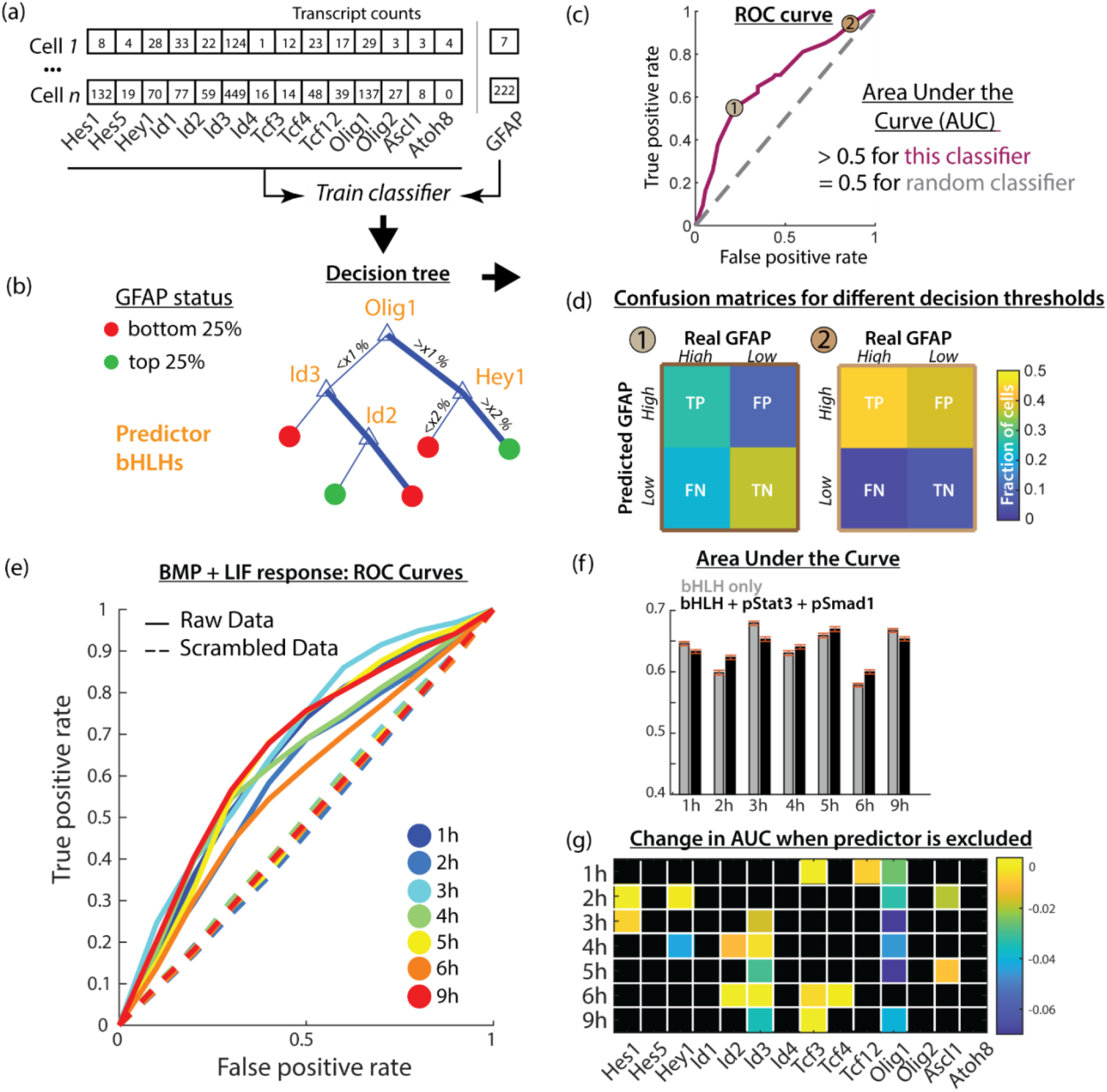
Expression levels of 14 bHLHs in a cell has predictive power about its GFAP status, with different sets of bHLHs dominating the decision at different times. (a-d) Example demonstrating classification of cells into GFAP low/high categories based on bHLH expression levels, and the key metrics used for assessment of predictive power. (a) For each cell, expression counts of 14 bHLH genes and GFAP (levels determined by sequential HCR RNA-FISH and corresponding to data shown in Figure S3) were used to train a decision tree classifier. (b) Example of a classification tree highlighting the bHLHs chosen as predictors after training and the decisions that result in cells being predicted to be GFAP-high or GFAP-low. Decisions are based on expression levels of predictors being above or below a percentile cutoff (e.g., x1, x2, etc.). Exact cutoff values depend on the decision threshold value, which determines prediction accuracy and recall (see (d)). (c) Predictive power of the classifier is captured in a Receiver Operating Characteristic (ROC) curve, showing how the True Positive Rate and False Positive Rate change relative to one another as the underlying decision threshold value is varied. A classifier with predictive power has an Area Under the Curve (AUC) larger than 0.5 since random classification would produce an AUC of 0.5. (d) Confusion matrices for two different decision threshold values, corresponding to 1 and 2 on the curve in (c). Each square shows the relative fraction of cells for true positive (TP), false positive (FP), true negative (TN) and false negative (FN) predictions. (e) Solid lines - ROC curves for classifiers trained on bHLH and GFAP expression levels (shown in Figure S3) at each of 7 different time points following treatment with BMP + LIF. Dashed lines - Corresponding ROC curves for classifiers trained on scrambled data. Note that classifiers trained on the raw data have higher AUCs than those trained on scrambled data, which behave like random classifiers. (f) Matrix indicating predictors (colored squares) chosen during classification training for each of the analysis timepoints (rows) following BMP + LIF treatment. The color of each predictor square corresponds to the contribution of the predictor to classifier Area Under the Curve (AUC).

Finally, we assessed the contribution of bHLHs chosen as predictors for each trained classifier by calculating the change in AUC when each specific predictor bHLH was excluded from the data used to train the corresponding classifier (Figure 3g). We observed that frequently used predictors like Olig1 and, to a lesser extent, Id3 had a bigger effect on the classifier performance compared to other predictor bHLHs, suggesting that they carry relatively more information about GFAP levels. Most importantly, no single bHLH was solely responsible for classification ability because AUC consistently remained ∼0.55-0.6 even when these bHLHs were excluded.

Together, these data show that the mRNA levels of 14 bHLHs in each cell have predictive power about its GFAP expression. At each timepoint following BMP + LIF treatment, this information is distributed across multiple bHLHs. Moreover, the set of bHLHs that contribute most to predictive power changes over time, suggesting a dynamic relationship between bHLHs and GFAP in the cell.

### LIF and BMP relieve repression of GFAP through a bHLH circuit

We next asked how bHLHs regulate GFAP expression. We first considered the Class I and Class II bHLHs Tcf4, Tcf12, Ascl1, and Olig2, which are downregulated upon BMP + LIF treatment (Figure 2c, Figure S2e). To test their role in GFAP expression, each of these bHLHs was ectopically expressed as an iRFP fusion under control of a doxycycline (dox) inducible promoter. Their effect on GFAP expression was assessed 24h after BMP + LIF + dox treatment (Figure 4a), allowing sufficient time for inducing ectopic bHLHs and expressing GFAP protein. Cells were binned based on their iRFP fluorescence, which reflected the level of ectopic bHLH expression, and the fraction of GFAP positive cells within each bin was calculated. This analysis showed that each of these bHLHs could suppress GFAP expression in a dose-dependent manner. On the other hand, knocking down different subsets of these genes using siRNA increased GFAP expression, but modestly (<2-fold, Figure S4a, b). These data show that multiple bHLHs can repress GFAP expression and their suppression is required to achieve strong GFAP expression.

**Figure 4:**
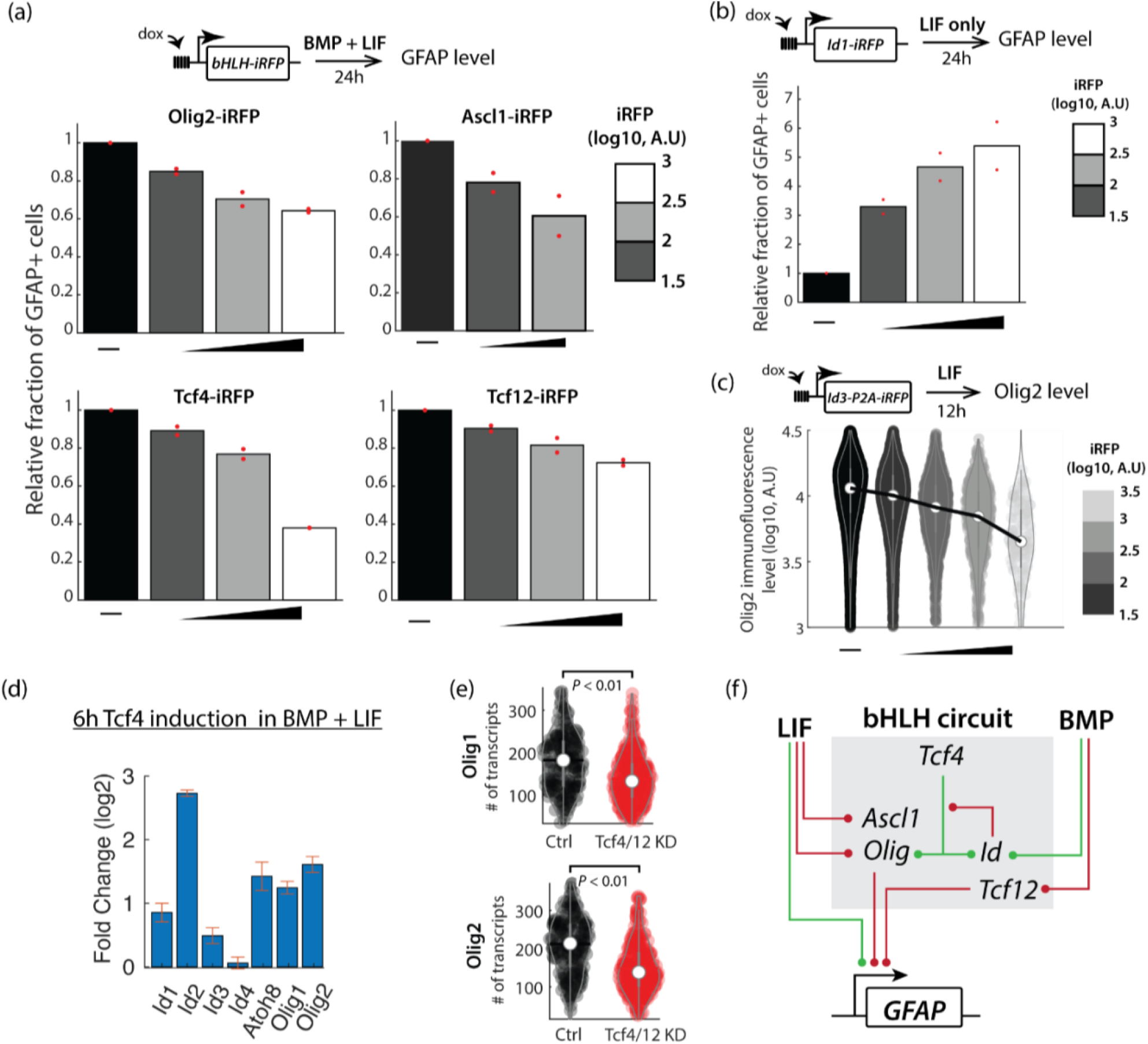
Downregulation of multiple repressive bHLHs by BMP and LIF is required for GFAP expression. (a) (*Top*) Experiment schematic: bHLH-iRFP fusion proteins were ectopically expressed using doxycycline (dox) at the same time as BMP + LIF treatment. GFAP levels were measured by immunofluorescence after 24h. (*Bottom*) Fraction of Olig2-iRFP, Ascl1-iRFP, Tcf4-iRFP, or Tcf12-iRFP expressing cells that were GFAP-positive, relative to cells expressing background iRFP levels (‘—’). Cells were binned based on their iRFP expression level. Red dots represent 2 biological replicates and the median is plotted. (b) (*Top*) Experiment schematic: Id1-iRFP fusion protein was ectopically expressed using doxycycline (dox) at the same time as LIF treatment. GFAP levels were measured by immunofluorescence after 24h. (*Bottom*) Fraction of iRFP positive cells that were GFAP-positive, relative to cells expressing background iRFP levels (‘—’). Cells were binned based on their iRFP expression level. Red dots represent 2 biological replicates and the median is plotted. (c) (*Top*) Experiment schematic: Id3-P2A-iRFP was ectopically expressed using doxycycline (dox) at the same time as LIF treatment. Olig2 levels were measured by immunofluorescence after 12h. (*Bottom*) Violinplots showing distribution of Olig2 levels in cells binned based on their iRFP level. Circles correspond to median Olig2 value. All pairwise comparisons of Olig2 distributions between bins are significant at a *P-*value < 0.01 by two-sided KS test. (d) Fold-change in the expression of the indicated genes, measured using RT-qPCR, after 6h of ectopic Tcf4-iRFP induction in the presence of BMP + LIF, compared to the no induction case. Error bars indicate S.E.M (n = 2 biological replicates). (e) Violinplots showing the number of Olig1 or Olig2 transcripts detected by HCR-FISH in cells that were treated with siRNAs targeting Tcf4 and Tcf12 (‘Tcf4/12 KD’) or control siRNA (‘Ctrl’) for 16h in growth conditions. *P*-value calculated using the two-sided KS-test (f) Schematic of the bHLH circuit based on data from this work including signal inputs from LIF and BMP, internal interactions, and GFAP-regulatory outputs. Green connections indicate positive interactions. Red connections indicate inhibition.

We next ascertained the role of the Id proteins in suppressing Class I and II bHLHs during astrocytic differentiation. Previous work has shown that Id proteins, induced by BMP (Figure 2c), can suppress the activity of Class I or Class II bHLHs through dimerization [14]. Indeed, we found that ectopic expression of Id1 or Id3 alone could upregulate GFAP expression in LIF, suggesting that the effects of BMP are mediated primarily through Id genes (Figure 4b, Figure S4c).

Our analysis unexpectedly revealed that ectopic Id3 expression leads to the downregulation of Olig1 and 2 at the mRNA level. This reduction was also apparent at the protein level for Olig2 (Figure 4c, Figure S4d). What underlies this effect? Since Id proteins do not bind directly to DNA, but inhibit the activity of other DNA-binding bHLHs such as the Tcf proteins, we asked whether the effect of Id on Olig transcription could be mediated through Tcf proteins. Indeed, ectopic expression of Tcf4 upregulated Olig1 and Olig2 expression within 6h (Figure 4d). Furthermore, knocking down Tcf4 and Tcf12 downregulated Olig1 and Olig2 mRNA levels in the absence of BMP + LIF (Figure 4e, Figure S4e). We suggest that the time required for BMP to relieve GFAP repression through this multi-step process (BMP induces Id, Id blocks Tcf to inhibit Olig expression, which leads to downregulation of Olig protein, finally relieving repression of GFAP) could explain the observed delay in the effect of BMP on GFAP transcription (Figure S1e).

We note that LIF also decreases Olig expression (Figure 2c) without inducing the Id genes. Therefore, LIF and BMP likely downregulate Olig genes through a separate mechanism, explaining their additive effects on Olig suppression (Figure S4f).

Together, these results suggest that multiple repressive bHLHs must be suppressed to allow maximal GFAP expression. To achieve this, LIF and BMP independently target a subset of bHLHs, like Ascl1 and Tcf12, while cooperatively target others, like Olig1 and Olig2, through a combination of Id-dependent and independent mechanisms. Thus, LIF and BMP operate through an interconnected bHLH circuit to regulate GFAP expression and the related cell fate.

## Discussion

A central question in stem cell biology is how cells integrate signaling information to choose between cell fate programs. Here, we find that in neural stem cells, the signaling ligands LIF and BMP synergistically induce an early marker of astrocyte differentiation GFAP (Figure 1) through a set of bHLH factors (Figure 2) that work together to ensure that GFAP is strongly derepressed only when both signals are present (Figure 4). Since the bHLH factors that must be suppressed to enable synergistic GFAP activation also promote alternative neuronal and oligodendrocytic fates, this system provides a tight coupling between signal integration and fate choice.

### Role of the circuit

An important feature of this system is that bHLH factors operate as a circuit, connected to each other through protein dimerization and transcriptional interactions. This contributes to signal integration in two ways: first, changes induced by BMP signaling enhance those initiated by LIF signaling. In particular, Ascl1 and Olig1/2, which promote neuronal and oligodendrocytic differentiation, respectively, are repressed by LIF and BMP through separate mechanisms. LIF transcriptionally downregulates their levels, whereas BMP induces the Id genes, which inhibit Ascl1 and Olig1/2 through heterodimerization. Second, circuit interactions enable even a single signaling pathway to regulate factors in multiple, mutually-reinforcing ways. For instance, Id proteins induced by BMP regulate Olig1/2 activity by (i) competing with Olig1/2 for dimerization with Tcf proteins, (ii) by blocking Tcf activity to decrease Olig1/2 transcription, and (iii) by directly binding with Olig proteins to generate inactive dimers [15].

### GFAP transcription dynamics

A surprising finding of this work was that GFAP transcription commences within 1h following treatment with LIF, with or without BMP (Figure 1). Differences in GFAP levels between signaling conditions primarily reflect differences in the temporal dynamics of GFAP. In particular, while LIF is sufficient to initiate GFAP transcription, BMP is required to maintain it. In fact, the contribution of BMP to GFAP expression has an inherent delay despite rapid activation of Smad1, the nuclear effector of BMP signaling (Figure S1e). We suggest that the delay reflects the time required for BMP, acting indirectly through Id factors, to decrease levels of Olig1 and Olig2, necessary for maximal GFAP expression.

### Relationship to Smad1-Stat3 cooperativity model

Previous work has ascribed synergy in GFAP transcription to an indirect cooperative interaction between Smad1, activated by BMP signaling, and Stat3, activated by LIF signaling [5]. This model is complicated by the absence of known Smad1 binding sites, particularly close to Stat3 sites, in the GFAP promoter [5,21,22]. While our work did not directly test the relative contribution of this interaction to GFAP regulation, our results suggest that this model is insufficient to explain synergy: (1) Ectopic expression of Id1 or Id3 (normally induced by BMP) can bypass BMP pathway activation to induce GFAP expression (Figure 4b, S4c). This result is consistent with previous work demonstrating that the effect of BMP on GFAP requires intermediate protein synthesis [22]. (2) Time delay in the effect of BMP on GFAP induction suggests a requirement for intermediate steps involving transcription and translation of Id proteins and degradation of Olig proteins (discussed above).

### The importance of transcriptional bHLH regulation

The conclusions of this work are supported by previous findings showing that Id3-Tcf3 dimerization promotes astrocyte differentiation in the brain following injury [23] and that Id-Olig dimerization inhibits oligodendrocytic differentiation [15]. More generally, however, while there has been much focus on regulation of bHLHs through dimerization, we find that transcriptional regulation also plays a key role in this system. BMP and LIF signaling regulated all analyzed bHLHs at the mRNA level. Tcf4 regulates the mRNA levels of multiple bHLHs in the circuit, like Ids, Atoh8 and Olig genes (Figure 4). Based on previous CHIP-seq analysis of Tcf4 binding sites, these likely reflect direct transcriptional control at least for some genes like Olig2 [24].

Our findings that multiple bHLHs can suppress astrocyte differentiation (Figure 4), that LIF and BMP signaling information is distributed across the entire bHLH circuit (Figure 2), and that different bHLHs carry information regarding GFAP status at different times (Figure 3) highlight the importance of studying bHLHs at the level of circuits. bHLHs are involved in fate regulation in a wide array of tissues [25]. Many of the bHLHs analyzed here, such as Id factors, Hes/Hey1 and Tcf factors are directly involved in numerous other neural and non-neural contexts, as are LIF and BMP signaling. We anticipate that the principles of signal integration and fate choice by bHLH circuits described here will therefore apply to many of these systems.

## Acknowledgements

We would like to thank Sean Megason and members of the Lahav and Megason labs for helpful discussions.

## Funding

Research in the Lahav Lab is supported by the National Institutes of Health grant R35 GM139572, and by the Ludwig Center at Harvard. N.N. is the Philip O’Bryan Montgomery, Jr., MD, Fellow of the Damon Runyon Cancer Research Foundation (DRG-2339-18).

## Competing interests

The authors declare no competing interests.

## Supplementary Materials

Materials and Methods

Supplementary Figures S1 to S4

## Notes

### Competing Interest Statement

The authors have declared no competing interest.

