## Supplementary Figures for "Signal integration by a bHLH circuit enables fate choice in neural stem cells"

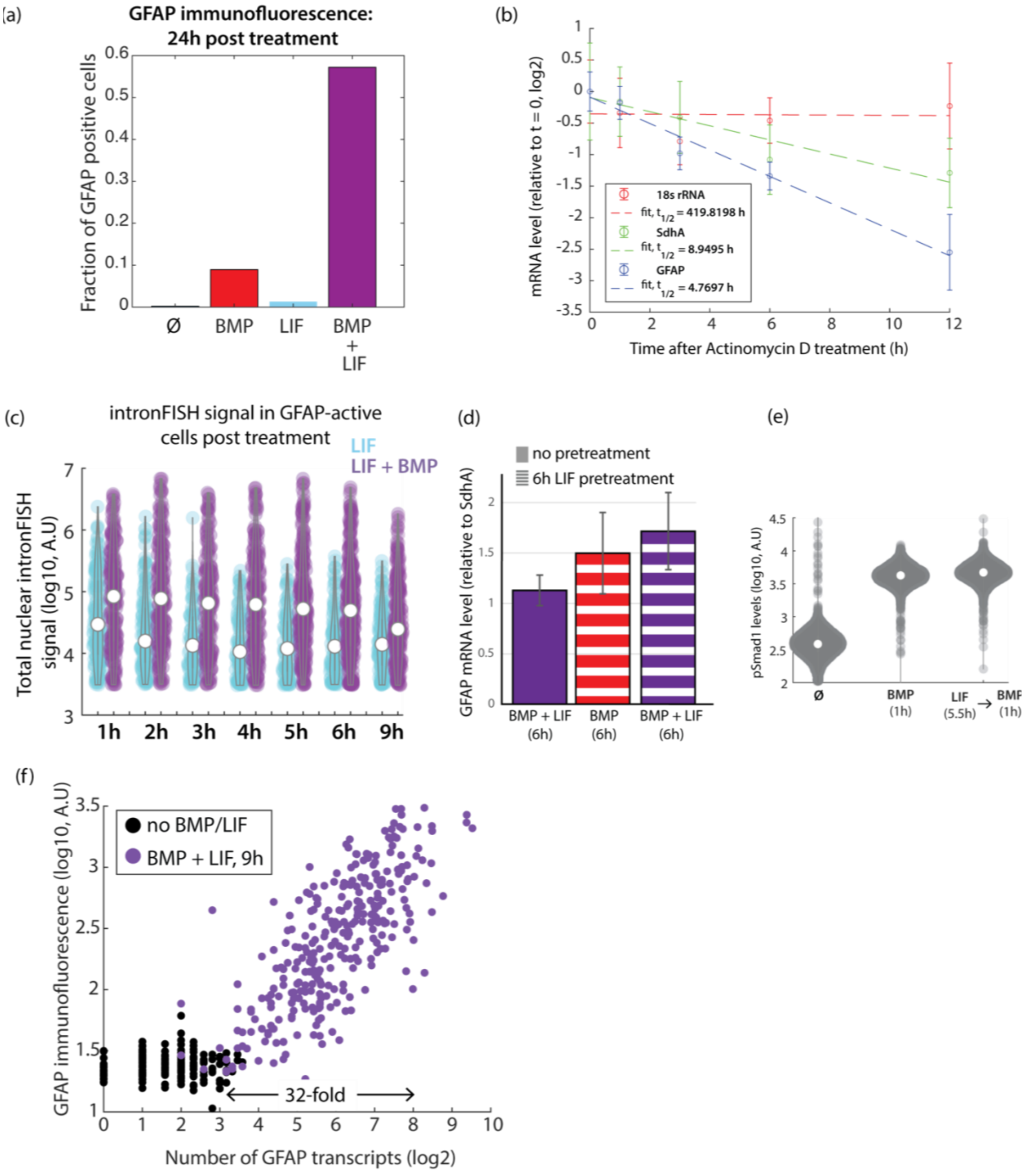

**Figure S1**: (a) Fraction of GFAP positive cells at 24h after treatment with BMP, LIF, BMP + LIF, or no treatment ('Ø'). (b) RT-qPCR quantification of mRNA levels (circles) following treatment with Actinomycin D, along with linear fits (dashed lines, see Methods). The half-life values ('$t_{1/2}$') corresponding to each fit are indicated. (c) Violinplots showing total intensity of the intronFISH signal in GFAP-active cells at the indicated times post treatment with LIF or BMP + LIF (same data as Figure 1e). The difference between LIF and BMP + LIF is significantly different at each timepoint (P-value < 0.01) by the two-sided KS-test. (d) RT-qPCR measurements of GFAP expression after 6h of combined BMP + LIF treatment (solid purple bar) or 6h of LIF pre-treatment followed by 6h of BMP or BMP + LIF treatment (red and purple hatched bars, respectively). Note that the LIF-pretreatment data is the same as that corresponding to the 6h timepoint in Figure 1e. (e) Violinplots showing phospho-Smad1 levels in untreated cells or cells treated with BMP for 1h, with or without LIF pre-treatment for 5.5h. (f) Scatter plot comparing number of GFAP transcripts detected using HCR RNA-FISH and GFAP immunofluorescence levels in cells treated with BMP + LIF for 9h (purple) or maintained in self-renewal conditions (black).

-

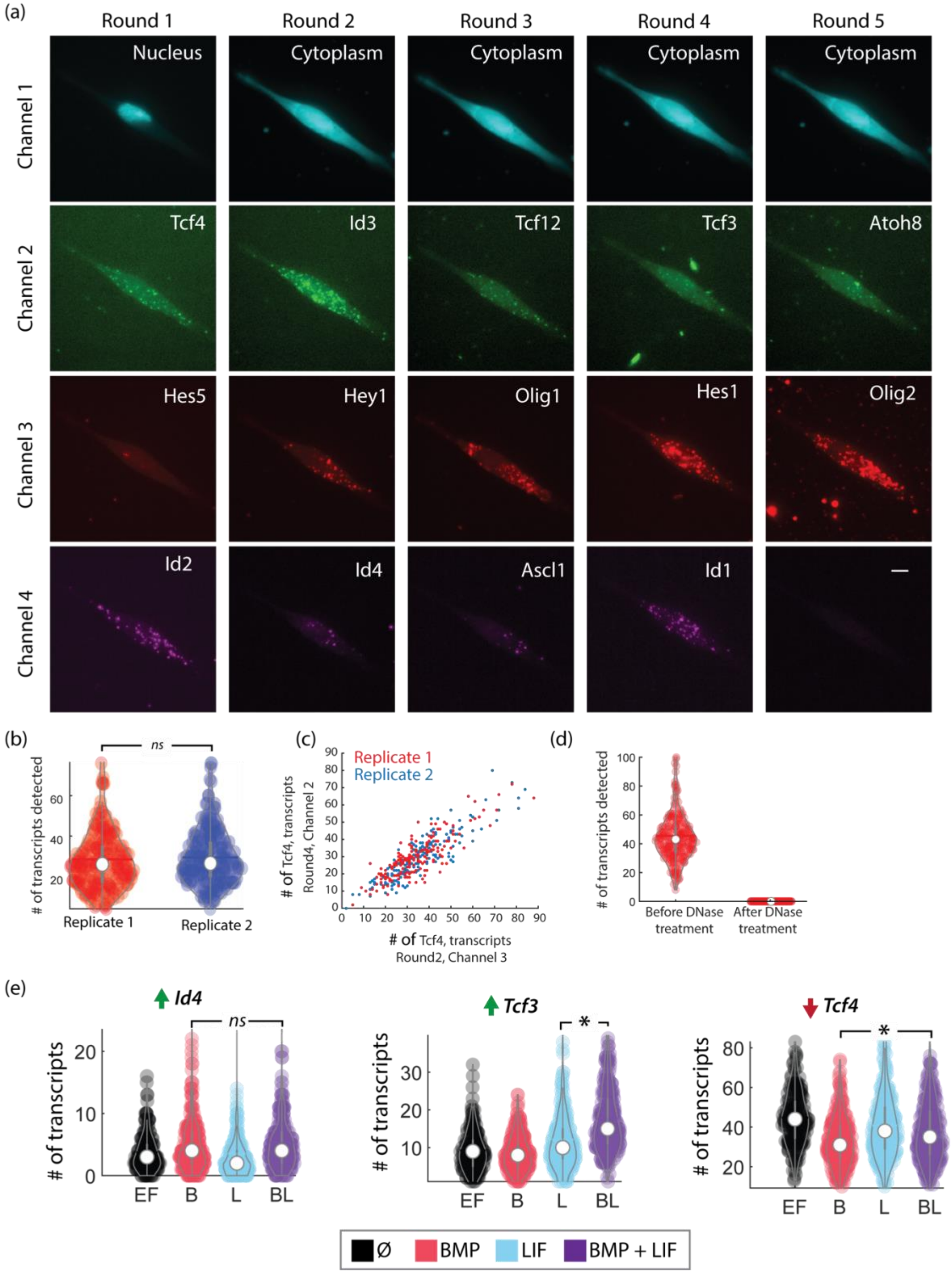
(a)
Round 1
Round 2
Round 3
Round 4
Round 5
Channel 1
Channel 2
Channel 3
Channel 4
Nucleus
Cytoplasm
Cytoplasm
Cytoplasm
Cytoplasm
Tcf4
Id3
Tcf12
Tcf3
Atoh8
Hes5
Hey1
Olig1
Hes1
Olig2
Id2
Id4
Ascl1
Id1
—
(b)
ns
### of transcripts detected
Replicate 1
Replicate 2
(c)
Replicate 1
Replicate 2
### of Tcf4, transcripts Round4, Channel 2
### of Tcf4, transcripts Round2, Channel 3
(d)
### of transcripts detected
Before DNase treatment
After DNase treatment
(e)
Id4
Tcf3
Tcf4
ns
*
*
### of transcripts
EF
B
L
BL
Ø
BMP
LIF
BMP + LIF

**Figure S2**: (a) Representative images of RNA-FISH signal for all 14 bHLHs in a single cell, measured across 5 rounds of sequential HCR RNA-FISH. Each image shows a maximum projection of 20 Z-slices. Different colors in the images correspond to different multiplexed spectral channels. Channel 1 was used to detect a nuclear marker in round 1 and cytoplasmic stain in subsequent rounds. (b – d) Quality controls for sequential HCR RNA-FISH. (b) Representative violinplots comparing number of mRNAs detected for a bHLH gene (Ascl1) in two biological replicates. *ns* indicates no significant difference in the distributions, by two-sided KS-test. (c) Representative comparison of mRNA transcript quantification for a bHLH gene (Tcf4) detected in channel 3 in round 2 of sequential HCR RNA-FISH and then detected again in round 4, in channel 2. This comparison is shown for two biological replicates (red and blue dots). Note that detection in round 4 corresponds to a fresh round of probe hybridization, since the probes used in round 2 were cleared before the subsequent round of RNA-FISH. (d) Representative data showing quantification of detected mRNA transcripts for a bHLH gene before and after the probe clearance using DNaseI treatment. (e) Number of mRNA transcripts detected per cell, using sequential HCR RNA-FISH, for 3 bHLHs in self-renewal conditions ('Ø') or after treatment with BMP, LIF, or BMP + LIF for 6 hours. These genes display < 1.5 fold change, relative to no treatment condition, in all of the signal treatments. *'*'* indicates a significant difference at a *P*-value of <0.01 by two-sided KS-test, while *ns* indicates no significant difference. Same experiment as in Figure 2c.

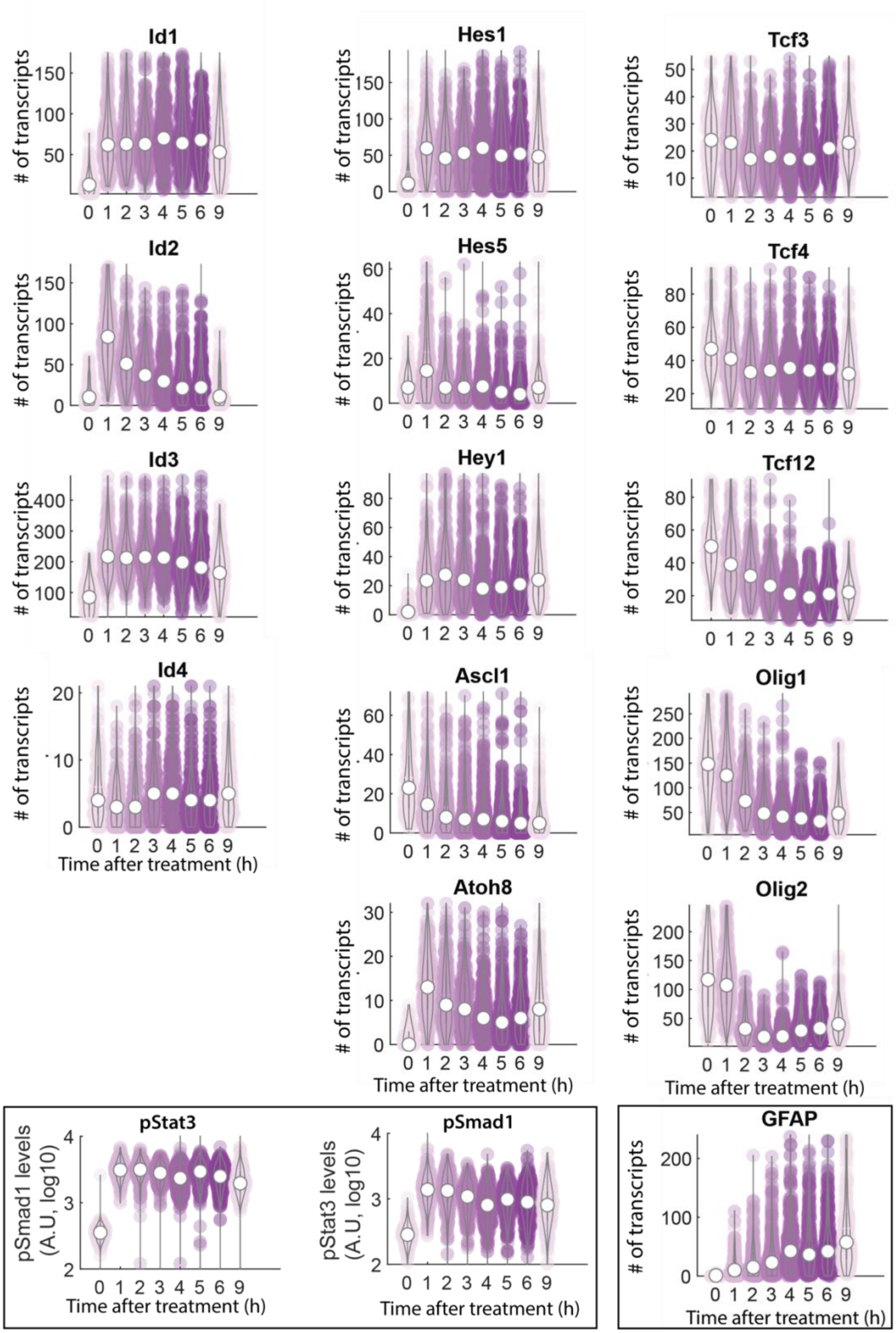


**Figure S3**: Violinplots showing mRNA transcript levels for bHLHs and GFAP, detected using sequential HCR RNA-FISH, as well as levels of phospho-Stat3 and phospho-Smad1, detected by immunostaining, during the first 9 hours after BMP + LIF treatment.

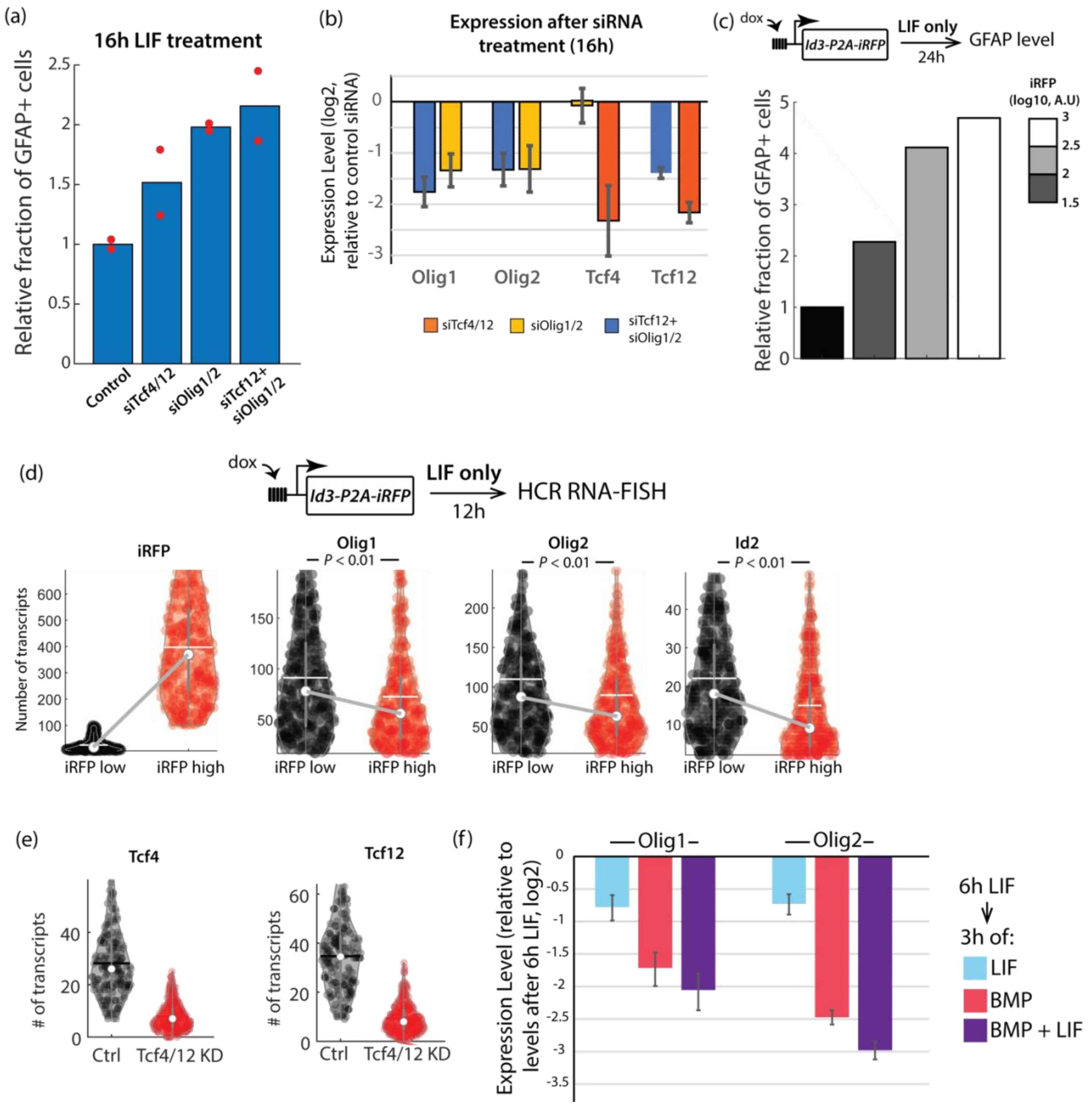

(a)
16h LIF treatment
Relative fraction of GFAP+ cells
Control
siTcf4/12
siOlig1/2
siTcf12+ siOlig1/2
(b)
Expression after siRNA treatment (16h)
Expression Level (log2, relative to control siRNA)
Olig1
Olig2
Tcf4
Tcf12
siTcf4/12
siOlig1/2
siTcf12+ siOlig1/2
(c)
dox
Id3-P2A-iRFP
LIF only
24h
GFAP level
Relative fraction of GFAP+ cells
iRFP (log10, A.U)
(d)
dox
Id3-P2A-iRFP
LIF only
12h
HCR RNA-FISH
iRFP
Olig1
Olig2
Id2
P < 0.01
Number of transcripts
iRFP low
iRFP high
(e)
Tcf4
Tcf12
### of transcripts
Ctrl
Tcf4/12 KD
(f)
Olig1
Olig2
Expression Level (relative to levels after 6h LIF, log2)
6h LIF
3h of:
LIF
BMP
BMP + LIF

**Figure S4**: (a) Relative fraction of GFAP positive cells 16h after treatment with LIF and siRNAs targeting the indicated genes (e.g., siTcf4/12 indicates treatment with siRNAs targeting Tcf4 and Tcf12). Red dots indicate two replicates and the median is plotted. Values are normalized to the mean GFAP fraction in Control siRNA-treated ('Ctrl') samples. (b) Expression levels of indicated genes after 16h of treatment with siRNAs in growth conditions. These siRNA-treated samples are the same as those used in (a). Error bars indicate S.E.M of two biological replicates. (c) (*Top*) Experiment schematic: Id3-P2A-iRFP was ectopically expressed using doxycycline (dox) at the same time as LIF treatment. GFAP levels were measured by immunofluorescence after 24h. (*Bottom*) Fraction of iRFP positive cells that were GFAP-positive, relative to cells expressing background iRFP levels ('—'). Cells were binned based on their iRFP expression level. (d) (*Top*) Experiment schematic: Id3-P2A-iRFP was ectopically expressed using doxycycline (dox) at the same time as LIF treatment. 12h later cells gene expression was measured using HCR RNA-FISH. (*Bottom*) Violinplots showing mRNA levels of the indicated genes in cells binned into 'iRFP high' and 'iRFP low' categories based on iRFP mRNA levels (leftmost panel). In each violinplot, circles indicate median, while white lines indicate mean value of the distribution. *P* values were calculated based on two-sided KS-test. (e) Violinplots showing the number of Tcf4 or Tcf12 transcripts detected by HCR-FISH in cells that were treated with siRNAs targeting Tcf4 and Tcf12 ('Tcf4/12 KD') or control siRNA ('Ctrl') for 16h in growth conditions. Same experiment as Figure 4e. (f) Relative expression level of Olig1 and Olig2, measured using RT-qPCR, in cells that were pre-treated with LIF for 6h and then treated for 3h with BMP, BMP + LIF or maintained in LIF. Expression levels shown were normalized by those at the end of the 6h LIF pre-treatment period.
