## Supplementary material for "Signal integration by a bHLH circuit enables fate choice in neural stem cells": Materials and Methods

### Neural stem cell culture and treatments

Cortical neural stem cells derived from E15-18 mice were obtained commercially (EMD Millipore, Catalog #SCR029). Cells were plated on tissue culture-treated plastic, coated overnight with poly-L-Ornithine (Millipore Sigma Catalog #P3655) and Laminin (Millipore Sigma Catalog #L2020) and maintained in neural basal medium (Millipore Sigma, Catalog #SCM003 or StemCell Technologies Catalog #05702) supplemented with 20 ng/ml recombinant EGF (Millipore Sigma Catalog #GF144), FGF-2 (Millipore Sigma Catalog #GF003) and Heparin (Millipore Sigma Catalog #H3149). Cultures were generally passaged using Accutase (Millipore Sigma Catalog #SF003) every 2 days or cryopreserved in liquid nitrogen in basal medium supplemented with 10% DMSO.

For LIF/BMP treatments, growth medium was replaced with neural basal medium typically containing 20 ng/ml recombinant BMP4 (R&D Systems, Catalog #314-BP) and/or 80 ng/ml recombinant LIF (R&D Systems, Catalog #8878-LF).

### Cell line construction

Most experiments were performed with a cell line that constitutively expressed a nuclear CFP marker and in which the Hes1 protein was endogenously tagged with an HA epitope tag. First, neural stem cells were co-transfected with CRISPR/Cas9 RNPs targeting the Rosa26 locus (guide RNA sequence: cgcccatcttctagaaagac) and an HDR (homology-directed repair) donor plasmid containing a mCerulean3-3xNLS gene downstream of a CAG promoter as well as a Neomycin resistance cassette. Antibiotic selection with 300 ug/ml G418 resulted in a polyclonal population in which the significant majority of cells were Cerulean-positive. Next, these cells were co-transfected with RNPs targeting the Hes1 gene (guide RNA sequence: gaaaatgccagctgatataa) and a single-stranded oligodeoxynucleotide (‘ssODN’, custom synthesized from IDT Technologies) designed to knock-in the HA epitope tag in-frame at the N-terminus of Hes1. A monoclonal cell line was isolated from this transfected population, verified by HA-staining and genomic sequencing, and used for further experiments.

For ectopic doxycycline-inducible expression of bHLH genes, cells were transfected with a piggybac plasmid vector containing the bHLH transgene (along with antibiotic resistance cassette) and a plasmid containing the super piggybac transposase. 24h after transfection, 10 ug/ml Blasticidin was added to select for cells in which the transgene had been stably integrated. Experiments were performed with the resulting polyclonal population or monoclonal lines isolated through limiting dilution.

### Cell transfection

For cell transfection, 1-2 x $10^6$ neural stem cells were typically electroporated using the Amaxa 2b Nucleofector (A-033 protocol) and nucleofection reagent kit (Lonza Biosciences, Catalog #VPG-1004) and plated onto poly-L-ornithine/Laminin-coated tissue culture plastic in growth medium.

### CRISPR/Cas9 targeting

Following the protocols from Dewari *et al* [1], CRISPR/Cas9 ribonucleoproteins (RNPs) were generated in vitro and nucleofected into cells for gene targeting. Briefly, synthetic tracrRNA (IDT Technologies Catalog #1072532) and gene-specific crRNA (custom synthesized from IDT Technologies) were first denatured and re-annealed together (1:1 ratio, 50 uM final concentration for each) and then mixed with 50 ug recombinant S.p Cas9 protein (IDT Technologies Catalog #1081058) at room temperature to generate RNPs.

**Plasmid construction**
Gibson cloning (using commercial reagents from NEB, Catalog #E2621) was generally used for plasmid construction.

HDR donor for constitutive Cerulean expression from Rosa26: An mCerulean3-3xNLS gene was cloned downstream of the CAG promoter and assembled into a plasmid vector along with with Rosa26 homology arms (from Addgene plasmid #61408) and a Neomycin resistance cassette.

piggybac-based vectors for ectopic bHLH expression: Each bHLH transgene was cloned downstream of a doxycycline-inducible TRE-tight promoter in a piggybac vector that also contained the rtTA-Advanced transactivator and Blasticidin resistance gene under control of constitutive promoters.

**Immunofluorescence assays**
For immunofluorescence assays, 3-5x$10^4$ cells were typically plated on 24-well glass-bottom plates (with 13 mm diameter wells) coated with poly-L-orninthine/Laminin. Following treatment, they were fixed in 4% Paraformaldehyde diluted in Phosphate-Buffered Saline (PBS), incubated in blocking buffer (PBS + 2% Bovine Serum Albumin + 0.3% Triton X-100) at room temperature and transferred to 4C overnight with primary antibody. After washes the next day they were incubated in 1:1000 Alexa-Fluor conjugated secondary antibodies either for 1h at room temperature or overnight at 4C.

The following primary antibodies were utilized: GFAP (Clone GA5, Cell Signaling Technology, Catalog #3670), Olig2 (Clone 2F11.1, Milllipore Sigma Catalog #MABN-50), mouse anti-pStat3 (Tyr705, Cell Signaling Technology Catalog #9145), rabbit anti-pStat3 (Tyr705, Cell Signaling Technology Catalog #4113), rabbit anti-pSmad1/5/9 (Cell Signaling Technology Catalog #13820).

**RT-qPCR**
RNA was extracted from cells using the RNeasy Mini Kit (Qiagen). 150-300 ng of RNA was reverse-transcribed into cDNA using the Multiscribe Reverse Transcription Kit (Applied Biosystems). 0.5 ul of cDNA was mixed with gene-specific primers (final concentration 450 uM) in Power Sybr Green Master Mix-based (Thermo Fisher Scientific) reaction. Real time amplification was analyzed on a CFX96 thermocycler (Biorad). The following primers were used for amplification:

| | |
|---|---|
| mAscl1 F | ACGACTTGAACTCTATGGCG |
| mAscl1 R | CAAAGTCCATTCCCAGGAGAG |
| mAtoh8 F | TCAACGGAGATCAAAGCCC |
| mAtoh8 R | AGTTTGGAGAGCTTCTGCC |
| mOlig1 F | AAGTTCCCGCATCTGGTC |
| mOlig1 R | GGAAGATTGGCTGAGGTCG |
| mOlig2 F | CGCAAGCTCTCCAAGATCG |

| | |
|---|---|
| mOlig2 R | CTCACCAGTCGCTTCATCTC |
| mHes1 F | GGCGAAGGGCAAGAATAAATG |
| mHes1 R | GTGCTTCACAGTCATTTCCAG |
| mHes5 F | CGGTGGAGATGCTCAGTC |
| mHes5 R | CTTGGAGTTGGGCTGGTG |
| mHey1 F | TACCCAGTGCCTTTGAGAAG |
| mHey1 R | TCCGATAGTCCATAGCCAGG |
| mId1 F | GCTGAACTCGGAGTCTGAAG |
| mId1 R | GCCTCAGCGACACAAGATG |
| mId2 F | CATCCCACTATCGTCAGCC |
| mId2 R | ATTCGACATAAGCTCAGAAGGG |
| mId3 F | GCATGGATGAGCTTCGATCTTA |
| mId3 R | CACCCAAGTTCAGTCCTTCTC |
| mId4 F | TGAACAAGCAGGGTGACAG |
| mId4 R | CGGTGGCTTGTTTCTCTTAATTTC |
| mTcf3 F | ATCTACTCCCCGGATCACTC |
| mTcf3 R | GGAGACCTGCATCGTAGTTG |
| mTcf4 F | CACAAACCATTACAGCACCTC |
| mTcf4 R | GTGTGGTCAGGAGAATAGATCG |
| mTcf12 F | AAGACCGCTCCATGATTCTG |
| mTcf12 R | TGGGAGATGGGTAACTAGGAG |
| mSdhA F | CCTACCCGATCACATACTGTTG |
| mSdhA R | AGTTGTCCTCTTCCATGTTCC |
| GFAP F | GAAAACCGCATCACCATTCC |
| GFAP R | CTTAATGACCTCACCATCCCG |
| 18s F | GAGACTCTGGCATGCTAACTAG |
| 18s R | GGACATCTAAGGGCATCACAG |

**Actinomycin D treatment and half-life estimation**
Cells were first treated with BMP + LIF for 24h to induce GFAP mRNA and subsequently treated with 10 µg/ml Actinomycin D (Biotium, Catalog # 10018-186). Cells were lysed for RNA extraction at the indicated times and analyzed by RT-qPCR.

To estimate the half life (‘$t_{1/2}$’) for each gene, a linear fit to its time-dependent change in Cq values (log2) relative to t = 0 was calculated. The inverse of the slope of this line corresponds to the $t_{1/2}$.

**Microscopy**
Samples were imaged in a Nikon widefield inverted epifluorescence microscope (Eclipse TE-2000) equipped with the Perfect Focus System (PFS) and LED light source (Xcite). Metamorph imaging software (Meta Imaging) was used for automatic image acquisition.

Immunofluorescence: A 20x (0.75 NA, air) objective was used for imaging immunofluorescence samples. Typically, ~150-200 fields of view were acquired for each sample by scanning across the dish.

RNA-FISH: A 60x (1.4 NA, oil) objective was used for imaging RNA-FISH samples. Z-stacks comprising 20 slices with 0.6 um spacing were acquired for each channel and field of view.

**HCR-FISH**
For each target gene, 16-20 DNA split-probes with hairpin toeholds were designed based on Choi *et al* [2] and synthesized using IDT Technologies. Fluorescently-labeled amplifier hairpins and buffers were purchased from Molecular Instruments. RNA detection was carried out based on protocols from the manufacturer and previous work [2,3]. RNAse-free reagents and plasticware was used throughout. Briefly, cells were fixed using 4% Paraformaldehyde (PFA) in Phosphate-Buffered Saline (PBS), then incubated with probes (in hybridization buffer) in a humidified chamber at 37C for approximately 24 hours. Following washes, bound probes were amplified for 1 hour at room temperature. Samples were stained using a 5-15 min incubation with CellTracker Violet dye (Thermo Fisher Scientific Catalog #C10094) during the wash steps before imaging.

For GFAP intronFISH experiments, probes were designed to target the first three introns of GFAP. The standard HCR-FISH protocol was followed for detection.

sequential HCR-FISH experiments consisted of up to five rounds of multiplexed HCR-FISH, where three genes were detected per round as detailed above. After microscopy to visualize mRNA transcripts, samples were treated with 1 U/ul DNAse I (Millipore Sigma Catalog #4716728001) for 3h at 37C, which clears signal and bound probes without affecting mRNA integrity. Degraded DNA was washed out with 20% formamide in 2x SSCT (Saline Sodium Citrate with 0.1% Tween-20) before proceeding to the hybridization step for the next round of HCR-FISH.

sequential HCR-FISH followed by immunostaining: Following the last round of HCR RNA-FISH, probes were cleared and samples were blocked in PBS + 0.3% Triton X-100 + 2 % BSA. The standard immunofluorescence protocol (see above) was followed subsequently.

**HCR-FISH signal quantification**
Custom MATLAB scripts were used to automatically segment cell nuclei and cytoplasm, detect HCR-FISH dots across multiple rounds and channels and assign them to individual cell segments.

Cell segmentation and registration: Nuclei and cell bodies in the image were segmented using a combination of edge-detection- or thresholding- based heuristics. Identifying nuclear segments enabled watershed-based splitting of conjoined cell body segments. Images of the same field of view across multiple rounds were registered based on cross-correlation in order to match cell and nuclear segments across rounds.

Dot detection: HCR-FISH signal appears as three-dimensional diffraction-limited spots or 'dots' in the imaging volume. Individual dots were identified as local maxima within voxels of localized signal that could be segmented from background based on intensity thresholds or edge-detection methods. While a threshold was chosen manually for each spectral channel based on visual inspection, the same thresholds were used across samples and HCR-FISH rounds for any given experiment.

MATLAB code used for analysis is available upon request.

**Immunofluorescence signal quantification**

For quantification of ectopic iRFP expression levels or fluorescent antibody staining levels, cell nuclei were segmented in images based on a constitutive nuclear marker (Cerulean-NLS) using standard edge-detection, thresholding and watershed analysis [4]. Fluorescence levels were quantified both within the nuclear segment and in a narrow annulus around each segment to estimate cytoplasmic signal levels. Median background fluorescence levels were subtracted from the median nuclear/cytoplasmic signal and used for analysis.

**Classification analysis**

The MATLAB ClassifierLearner app was used to train decision trees. First, to generate the training dataset, bHLH expression data was converted into Z-scores while GFAP expression was categorized into GFAP-high (top quartile of the distribution), GFAP-low (bottom-quartile) or GFAP-med (remaining data). Next, bHLH Z-scores in GFAP-high and GFAP-low cells were provided as predictor data to the classifier, along with the GFAP category. Finally, decision trees with a maximum of four splits were trained with fivefold cross-validation. To calculate median ROC curves, AUCs and bootstrapped error, this training procedure was carried out 100 times. To calculate ROC curves for scrambled data, the bHLH Z-score matrix was shuffled and trained with the same parameters and GFAP categories 100 times. Medians of the resulting ROC curves are shown in Figure 3e.

MATLAB code used for analysis is available upon request.

**Ectopic bHLH induction**

To induce expression of bHLH transgenes ectopically, 5-10 ug/ml doxycycline was added to culture medium.

**siRNA-based knockdown**

For knockdown experiments, 1-2x$10^6$ cells were transfected with 50 uM DsiRNA (IDT Technologies). Pre-designed DsiRNAs were used to target Olig1, Olig2, Tcf4 and Tcf12. For each experiment, a control transfection was performed with negative control DsiRNA (IDT Technologies Catalog #51-01-14-03).

| | |
|---|---|
| mm.Ri.Olig1.13.1-SEQ1: | rCrUrUrCrCrCrUrArArArGrGrUrArGrCrUrUrArArCrCrAAC |
| mm.Ri.Olig1.13.1-SEQ2: | rGrUrUrGrGrUrUrArArGrCrUrArCrCrUrUrUrArGrGrGrArArGrUrG |
| mm.Ri.Tcf12.13.1-SEQ1: | rUrUrCrArArGrGrUrArUrUrArUrUrGrArUrArArArCrCrUTT |
| mm.Ri.Tcf12.13.1-SEQ2: | rArArArGrGrUrUrUrArUrCrArArUrArArUrArCrCrUrUrGrArArUrU |
| mm.Ri.Olig2.13.1-SEQ1: | rGrCrArArGrCrUrUrUrArCrArGrArCrGrCrUrUrArArArATA |
| mm.Ri.Olig2.13.1-SEQ2: | rUrArUrUrUrUrArArGrCrGrUrCrUrGrUrArArArGrCrUrUrGrCrUrC |
| mm.Ri.Tcf4.13.1-SEQ1: | rGrUrGrUrUrUrCrUrArArUrUrArCrCrGrGrArUrArUrUrGAA |
| mm.Ri.Tcf4.13.1-SEQ2: | rUrUrCrArArUrArUrCrCrGrGrUrArArUrUrArGrArArArCrArCrUrA |
